# Quantitative muscle color as a proxy for structural and functional characteristics during muscle remodeling in *Gryllus lineaticeps*

**DOI:** 10.64898/2026.01.29.702701

**Authors:** Meghan Laturney, Lourenço Martins, Tomás Diaz, Evan Lo, Nio Uen, Caroline M. Williams

## Abstract

Understanding the cellular and physiological mechanisms underlying muscle remodeling requires model systems that allow rapid, reliable, and quantitative assessment of muscle state. The cricket *Gryllus lineaticeps* naturally undergoes non-pathological striated muscle breakdown (histolysis), making it a promising system for studying this process. However, current assessments of muscle state are largely qualitative, subjective, and poorly standardized across experiments. Here, we developed and validated a continuous, quantitative muscle color metric to objectively capture histolysis progression and functional changes in muscle. We show that this metric robustly tracks variation in muscle color across remodeling stages, including the challenging fully transparent stage, and strongly predicts protein content, mitochondrial abundance, and iron content in a muscle- and trait-specific manner. The reproducibility of these relationships across independent datasets demonstrates the generality and robustness of this approach. By providing a rapid, objective, and biologically informative proxy of muscle state, this framework not only advances the utility of *G. lineaticeps* as a model for muscle remodeling but also offers a strategy for exploring the cellular dynamics underlying age-related muscle diseases and disorders, addressing an increasing public health concern in aging populations.

## INTRODUCTION

Age-related muscle diseases and disorders are increasing worldwide and represent a growing public health concern in aging populations (Liu et al. 2019; Ru et al. 2025). Developing effective interventions requires a deeper understanding of the cellular and physiological pathways that govern muscle maintenance, remodeling, and resilience. An emerging model system for these processes is the California variable field cricket, *Gryllus lineaticeps*, which undergoes a natural, non-pathological form of striated muscle remodeling (Diaz et al. 2024, Treidel et al. 2023). Importantly, the intracellular mechanisms involved in cricket flight-muscle degeneration, such as altered autophagy-mediated protein turnover and mitochondrial abundance (Diaz et al. 2024) parallel those implicated in human myopathies (Arevalo et al. 2025, Wiedmer et al. 2021). However, a key limitation of this model is the subjective and poorly standardized nature of current methods for quickly assessing muscle functional state. Advancing the utility of this system, and its translational relevance, therefore requires the development of a rapid, accurate, and objective approach for quantifying muscle function. Such an approach would also be broadly applicable to other insect models of muscle function and dysfunction, including *Drosophila* or locusts (Protasiewicz et al. 2025, Guo et al. 2021).

*G. lineaticeps* crickets selectively breakdown their flight muscles, which are large, metabolically active tissues analogous to mammalian striated muscles (Ellington 1985, Iwamoto 2011). Cricket flight musculature includes the dorsal longitudinal flight muscles (DLMs) and the adjacent dorsal ventral muscles (DVMs). The DLM is used exclusively for flight and undergoes complete degeneration, whereas the DVM functions in both flight and walking and exhibits substantially less breakdown (Figure 1), providing a powerful within-animal control. This natural system enables precise investigation of healthy muscle remodeling, in which degradation, nutrient recycling, and protection from oxidative damage must be tightly coordinated.

**Figure 1.**
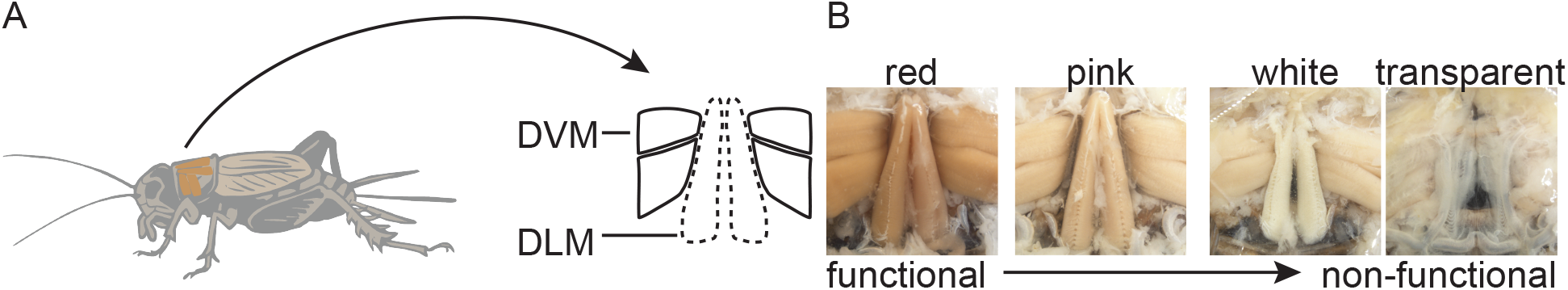
Flight muscle breakdown in *Gryllus lineaticeps*. A) The dorsal longitudinal muscle (DLM, outlined with a dashed line) is used for flight and dorsal ventral muscle (DVM, outlined with a solid line) is used for flight and walking. B) Images show stages of a continuous process of muscle breakdown from fully functional (left) to non-functional (right).

To rapidly assess flight muscle functional state, researchers commonly use muscle color as a proxy of metabolic capacity. Color categories are widely employed across insect flight muscle research, particularly in work on wing-dimorphic crickets, because muscle color is thought to reflect underlying functional properties. Darker, redder muscles typically indicate higher mitochondrial abundance, greater oxidative capacity, higher protein concentration, and potentially increased iron content, whereas paler muscles reflect reduced metabolic activity. (Diaz et al. 2024, Zera et al. 1997, Lu et al. 2023, Lorenz 2007). These relationships are well established in vertebrate physiology and are consistent with insect muscle biochemistry (Cao and Jin 2020, Iwamoto 2011).

Despite the utility of this model, assigning discrete color categories is inherently limiting. This framework imposes categorical boundaries on what is, in reality, a continuous physiological transition. Most studies distinguish only between “pink” muscles, considered functional, and “white” muscles, considered non-functional (Treidel et al. 2023, Clark et al. 2015, Zera et al. 1997). More recent work has refined this scheme by recognizing a third, transitional state: fully intact and functional muscles were labeled “red”, muscles undergoing degradation were designated as “pink” and fully histolyzed, non-functional, muscles were classified as “white” (Diaz et al. 2024). This revision reflects evidence that red (fully functional) and pink (histolyzing) muscles differ substantially in structure and function, and that in some cases, pink muscles more closely resembled white than red muscles (Diaz et al. 2024). Thus, the category formally labeled “pink” encompasses a heterogeneous set of muscle states, including tissues that are already partially or largely degraded, underscoring the need for a more refined and quantitative assessment of color. Moreover, the categorical approach relies on a subjective visual assessment of muscle color and lacks clear operational definitions, leading to poor standardization across studies.

The relationship between visible color and functional measures has never been quantitatively tested in *G. lineaticeps*, in large part because no objective metric of color exists. Here, we developed and validated a protocol for quantifying muscle color using standardized imaging and computational analysis. Our goal is to establish an unbiased, reproducible, and transferable method that improves accuracy within and across research groups. We then apply this quantitative color metric to evaluate its relationship with protein content, mitochondrial abundance, and iron content, key indicators of muscle functional state. Together, this protocol and validation provide the first objective framework for linking muscle color to muscle function in an established model of healthy striated-muscle remodeling.

## METHODS

### Cricket stocks

All experiments used adult wing-dimorphic crickets (*Gryllus lineaticeps*). The laboratory colony was founded in 2015 from adults (>200) collected from Sedgwick Reserve (34°41’34”N, 120°02’26”W, Santa Ynez, California). The colony is maintained at a constant size of 120 reproductively mature adults, with equal proportion of long-winged and short-winged crickets (50:50 wing morph ratio). To maintain genetic diversity, offspring from ∼100 field collected adults are added annually. Experiments used crickets from this lab colony unless explicitly stated.

All crickets used in experiments were maintained under the same environmental conditions (27°C, 16:8 h light:dark cycle; *ad libitum* access to water, and a standard laboratory diet, Treidel et al. 2023). Before adulthood, crickets were reared in clear plastic bins (60 cm x 35 cm x 45 cm) and sexes were separated at the last juvenile stage. To obtain virgin females of known age, bins were checked every 24 hours, and new adult females were housed individually in plastic pint containers.

### Development of color metric

Flight muscle color is commonly used as a qualitative proxy for muscle function, with red muscle indicating functional tissue and white muscle indicating non-functional tissue. However, this scoring is subjective, particularly when muscles are in intermediate stages of breakdown, and prone to variation among observers and laboratories. To reduce bias and improve reproducibility, we developed a digital method to quantitatively measure muscle color and objectively track the progression of histolysis.

### Training dataset preparation

We collected 67 long-winged and 36 short-winged females aged 3 to 7 days (n= 5-19 per day) to capture a broad range of muscle colors. Heads and legs were removed, and the thorax and abdomen were opened using fine-point micro-scissors. After removing the gut tract, the dorsal longitudinal muscles (DLMs) and the dorsal ventral muscles (DVMs) were exposed and manually categorized as red (R), pink (P), white (W), or transparent (T). Muscles with the darkest pigmentation were classified as red; those retaining visible pigmentation but substantially lighter in appearance were labeled as pink; muscles lacking visible pigment were labeled white; and tissues appearing clear were classified as transparent. Muscles were imaged using a Nikon SMZ18 stereomicroscope (Nikon Instruments, Melville, NY) and NIS-Elements software. To reduce glare and improve contrast, 50 μL of PBS was applied directly to the muscle. Images were saved as .tif files for subsequent digital analysis in ImageJ/FIJI.

### Digital muscle color quantification

Each multi-channel .tif image was separated into its constituent color channels (red, green, and blue) in ImageJ/FIJI, producing three grayscale images for independent analysis. Using the “Record” tool in the Macros plugin, we selected three non-overlapping regions of interest (ROIs) within each muscle type and recorded the mean pixel intensity for each ROI across all three channels, yielding nine measurements per each muscle type (Figure 2). These values were exported as .csv files and combined in Python using a JupyterLab Notebook (version 3.6.3).

**Figure 2.**
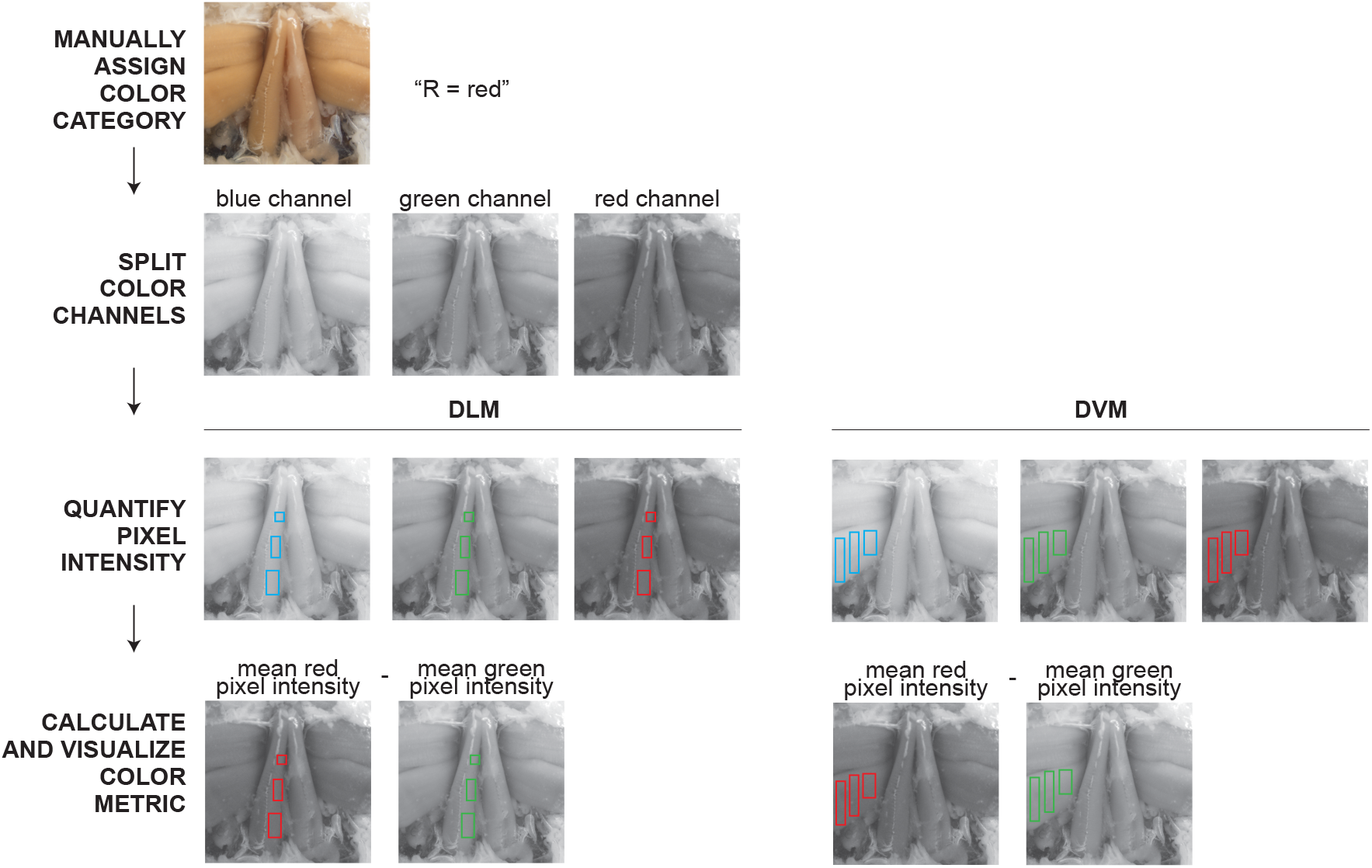
Workflow for muscle color metric development in the training dataset. Flow chart shows an example for one image. Manual muscle color category was determined by an observer (R = red, P = pink, W = white, T = transparent). Color channel separation and pixel intensity was performed in ImageJ. Color metric was calculated using the difference between the average red color channel value minus the average green color channel value.

### Exploring the relationship of muscle color and muscle function using mitochondrial abundance and iron content metrics

After developing the color metric, we explored the relationship between muscle color and two functional characteristics: mitochondrial abundance (citrate synthase assay) and iron content in two independent datasets. To link these datasets, we measured protein content in both.

### Test dataset #1: Measuring protein content and mitochondrial abundance

#### Test dataset #1 sample preparation

We collected 40 long-winged females on day 0 of adulthood, and aged individuals from 0 to 18 days to ensure a wide range of muscle colors and functional states. Each cricket was dissected at its assigned age, and the DLM and the DVM were isolated, weighed, imaged, and the color metric was calculated as described above. Following imaging, the muscle tissues were homogenized using a mortar and pestle in homogenization buffer (5 mM EDTA dihydrate, 50 mM of HEPES, and 0.1% Triton-X100) with a buffer volume equal to 25x the wet muscle mass.

Homogenates were centrifuged (10,000 x g, 2 min, 4ºC) and the resulting supernatant was collected and transferred to 2 mL microcentrifuge tubes. DVM samples were further diluted fivefold with homogenization buffer prior to downstream assays.

#### BCA Protein Quantification

Total protein concentration was quantified using the Thermo Scientific™ Pierce™ Dilution-Free™ Rapid Gold BCA Protein Assay Kit according to the manufacturer’s instructions (Thermo Fisher Scientific, Rockford, IL, USA). Protein standards were prepared using bovine serum albumin spanning a concentration range of 0-2 mg.mL^-1^. BCA assay reagent was prepared at a 50:1 ratio of Reagent A to Reagent B. For each sample and standard, 10 µL was pipetted in triplicate into a 96 well plate, followed by 200 µL of BCA assay reagent. Plates were incubated at room temperature (22-23 ºC) for 5 minutes and absorbance was measured at 480 nm using a SYNERGY H1 microplate reader.

#### Citrate synthase assay

Citrate synthase (CS) is a mitochondrial matrix enzyme that catalyzes the first step of the tricarboxylic acid (TCA) cycle and is a good proxy for mitochondrial content of muscles (Larsen et al. 2012). CS activity correlates well with mitochondrial function and whole organism aerobic performance in Gryllus crickets (Treidel et al. 2021). CS activity was measured spectrophotometrically by coupling CoA-SH production to a colorimetric reaction, providing an estimate of mitochondrial content in the tissue. For each sample, 10 µL was pipetted in triplicate into a 96-well plate.

Assay buffer (200 µL per well) containing 12 mM acetyl-CoA, 2.0 mM DTNB in 95% ethanol, and 50 mM Tris-HCl (pH 8.0) was added, and absorbance at 412 nm was recorded for 10 min at room temperature using a Synergy H1 microplate reader.

Subsequently, 5 µL of 21.5 mM oxaloacetate was added to each well, and absorbance was measured for an additional 10 min.

CS was calculated via the following equation:

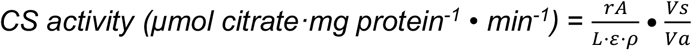

Where:

- rA is the rate of absorbance change (OD*min^-1^)
- L is the optical path length (cm)
- ε is the extinction coefficient (13.6 OD mM^-1^*cm^-1^)
- ρ is the protein concentration of the homogenate (mg.ml^-1^)
- Vis the total volume of the assay solution in each well (mL)
- Vis the volume of homogenate in the well (mL)

### Test dataset #2: Measuring protein and iron content

#### Test dataset #2 sample preparation

We collected 38 long-winged females on day 0 of adulthood and aged individuals 5 days, at which age approximately half of all females have initiated histolysis of their flight muscles (Treidel et al. 2023). Each cricket was dissected, and the DLM and the DVM were isolated, weighed, imaged, and the color metric was calculated as described above. Following imaging, the muscle tissues were flash-frozen in liquid nitrogen and stored at -80 ºC until use (up to 3 weeks). To prepare samples for subsequent assays, we thawed and homogenized tissue in 300 µl of lysis buffer (1% Triton X100, 1% SDS, 1X TBS, 1 mM EDTA, and 1% Protease inhibitor cocktail) per 5 mg of muscle tissue using a plastic pestle and then sonicated each sample with a handheld sonicator (1 x10s; Model 50, Fisher Scientific, Hampton, NH, USA). After sonication, samples were centrifuged (15,000 x g, 10 min, 4 °C), and the resulting supernatant was collected and transferred to 2 mL microcentrifuge tubes. Total protein was quantified using a BCA assay as described above.

#### Ferrozine assay

A ferrozine assay was performed as previously described (Soltani et al. 2024). For each sample, 77 µL of lysate was combined with 17 µL of concentrated HCl and incubated at 95 °C for 20 min to release protein bound iron. Following incubation, 50 µl of the boiled sample was mixed with 20 µl of 75 mM ascorbate (Sigma-Aldrich, #A92902-100G), vortexed, and centrifuged briefly. Next, 10 µL of 10 mM ferrozine (Sigma-Aldrich, #160601-1G) was added to each sample, followed by vortexing and centrifugation. Finally, 10 µL of 5 M ammonium acetate was added and samples were vortexed. A standard curve was generated using ammonium iron (II) sulfate (Sigma-Aldrich, #09719-50G) at concentrations ranging from 0 to 400 µM. Samples and standards were loaded in duplicate into a 96-well plate, followed by the addition of 50 µl of 10 mM ferrozine. Absorbance at 562 nm was measured using a Synergy H1 microplate reader.

## Statistical Analysis

To characterize the relationships between the color metric and three functional metrics (protein content, citrate synthase activity, and iron content), we evaluated three candidate models: linear, quadratic, and exponential. Model selection was based on Akaike’s Information Criterion (AIC), with the best-fitting model defined as the model with the lowest AIC value. Goodness of fit was assessed using the coefficient of determination (*R*^2^). Residual diagnostics were examined to verify that model assumptions were satisfied. To quantify the strength and direction of relationship between metrics, we performed a Pearson’s correlation on raw or log–log–transformed data for linear or exponential relationships, respectively.

## RESULTS

### Color metric captures variation in muscle color

The training dataset captured the full range of muscle phenotypes observed during flight-muscle histolysis, including red, pink, white, and transparent (fully histolyzed) muscles, providing a broad spectrum of coloration against which to evaluate quantitative color metrics. We first examined the raw pixel intensities of the red, green, and blue channels extracted from each image. Across manually assigned categories, the three channels showed a similar trend, with darker (red) muscles exhibiting lower mean pixel values and lighter (white) muscles exhibiting higher values (Figure 3A).

**Figure 3.**
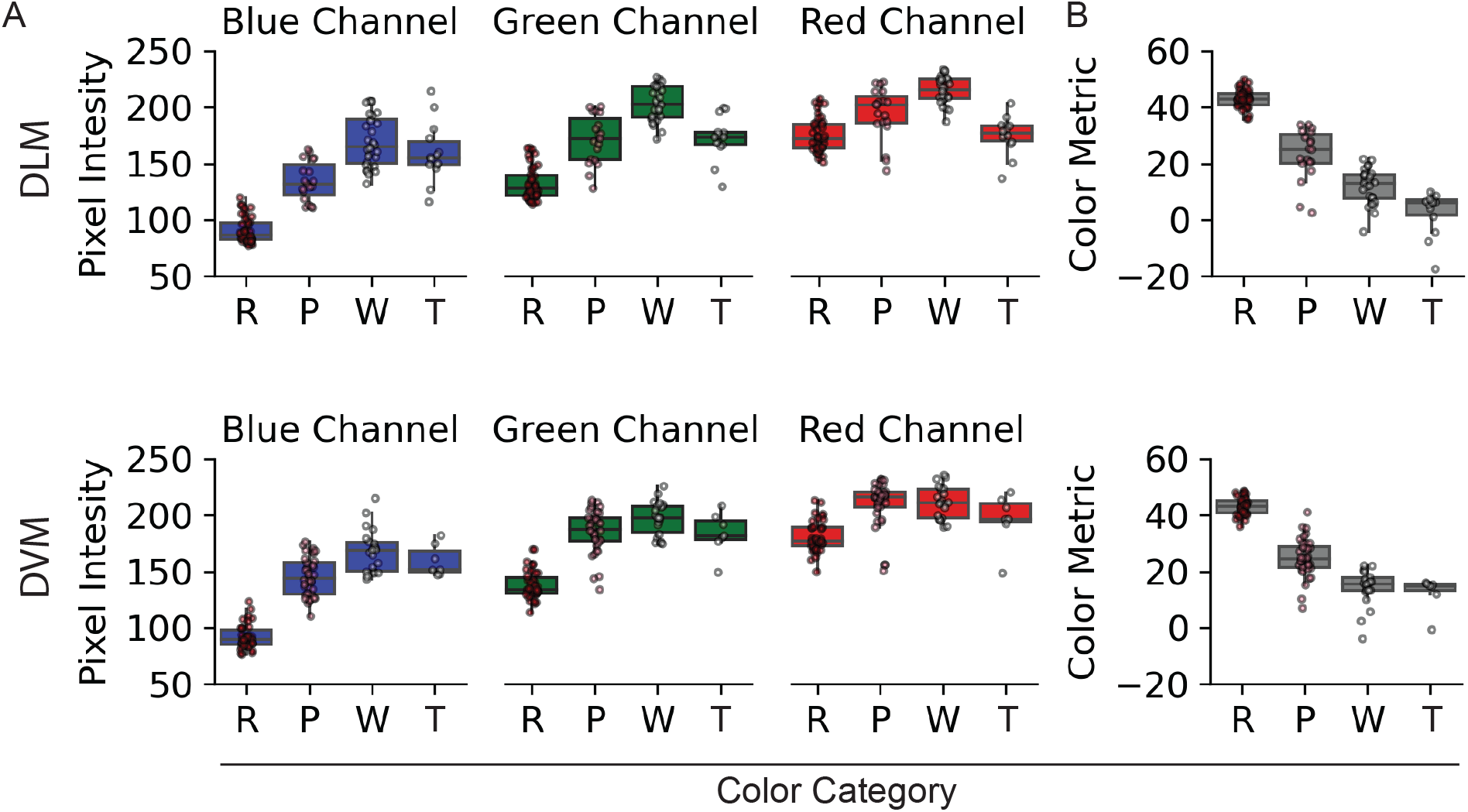
Color metric is a quantified value of muscle color. A) Average pixel intensity in the blue, green, and red color channels and B) the calculated Color Metric across manually assigned muscle color categories (R = red, P = pink, W = white, T = transparent) in DLM (top) and DVM (bottom) training dataset. Color Metric is calculated for each individual (average red channel pixel intensity - average green channel pixel intensity). Raw data is plotted as points, boxplots shows the quartiles, with the lower and upper edges of the box corresponding to the 25th and 75th percentile, and the whiskers indicate the values within +/-1.5 interquartile range (Tukey’s convention) to help identify extreme outliers.

However, transparent (fully histolyzed) muscles (the ‘T’ category; 14 DLMS and 7 DVMs) deviated from this pattern. Because these muscles were transparent, their pixel intensities were artificially low (and overlapping with the pink category) likely reflecting the color of the underlying cuticle coloration rather than the muscle itself (Figure 3A).

To develop a metric that better captured the full extent and progressive nature of histolysis, we evaluated combinations of color channels that might better distinguish true muscle coloration from background interference. A derived color metric, defined as the red-channel value minus the green-channel value, showed the strongest association with the manual labels, with higher values corresponding to darker red (more functional) muscle coloration. See Figure 3.

A simple difference metric, calculated as the red-channel value minus the green-channel value, provided the clearest separation and correct ordering of muscle categories (Figure 3B). This metric reduced the influence of transparency-associated background coloration and produced values that increased reliably with darker, more functional muscle. We therefore selected this difference score as our quantitative color metric for all subsequent analyses.

### Muscle color metric is related to protein content and mitochondrial abundance

We next evaluated whether the color metric predicts functional muscle properties, specifically protein content and mitochondrial abundance. Because short-winged females (which are not flight capable) do not use their dorsal longitudinal flight muscles (DLM), we restricted this analysis to long-winged females to validate the color metric against functional traits. We generated an independent test dataset consisting of 40 long-winged females with muscle colors spanning red, pink, and white categories. No crickets in this dataset had fully histolyzed, transparent muscles as all females were long-winged and transparent muscles are rare in this wing morph. As in the training dataset, the color metric captured substantial variation both within and among color categories, validating its performance in an independent sample not used for metric development (Figure 4A).

**Figure 4.**
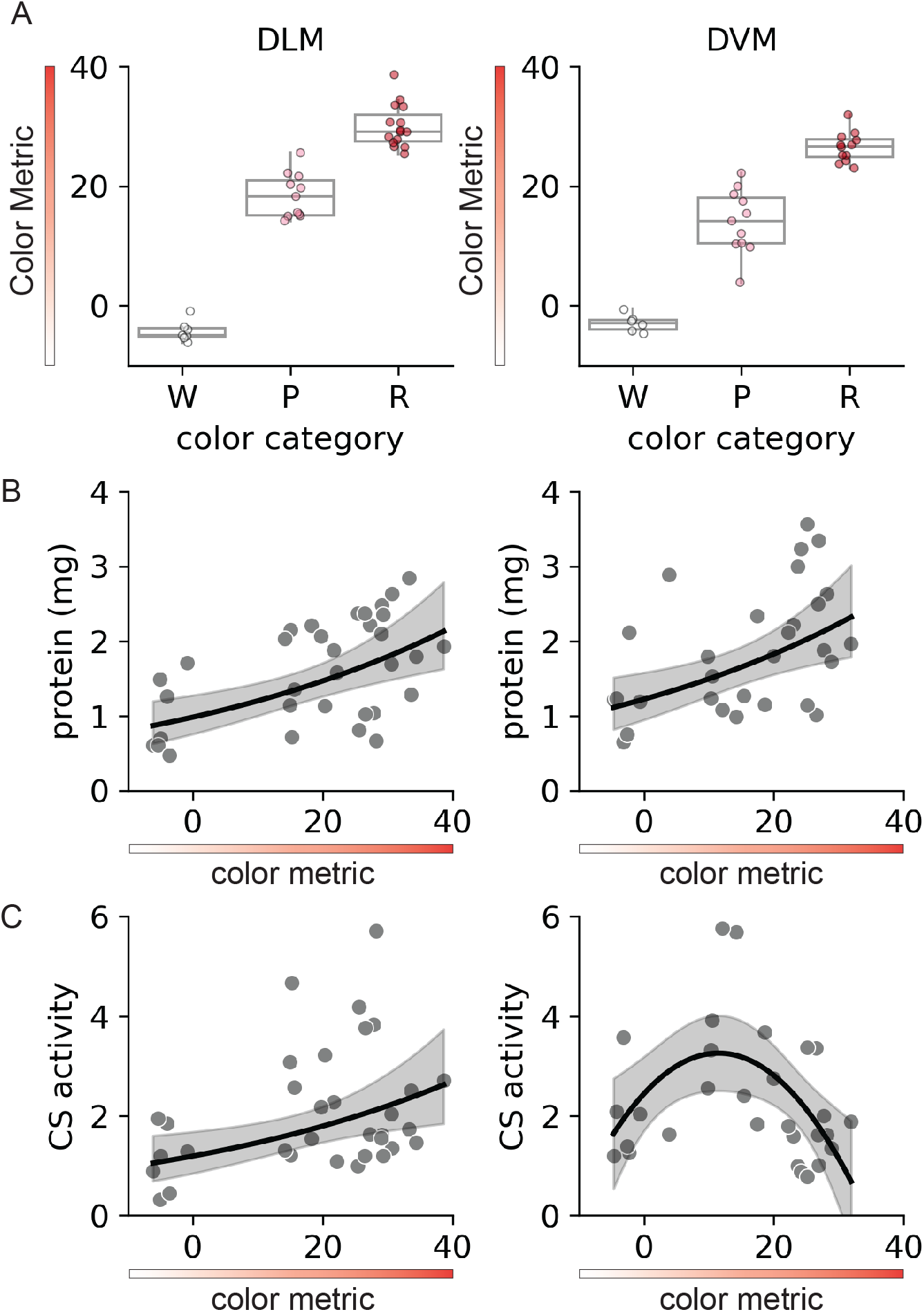
Muscle color is associated with protein content and mitchondrial abundance in both the DLM and DVM. A) Color Metric across manual color categories (R = red, P = pink, and W = white) in the DLM (left) and DVM (right). Raw data is plotted as points, boxplots shows the quartiles, with the lower and upper edges of the box corresponding to the 25th and 75th percentile, and the whiskers indicate the values within +/-1.5 interquartile range (Tukey’s convention) to help identify extreme outliers. Scatterplots showing the relatioship between color metric and (B) protein content and (C) citrate synthase (CS) activity, a proxy of mitochondrial abundance, in the DLM (left) and DVM (right). Solid lines represent the fitted exponential or quadratic model with shaded 95% confidence intervals.

We modeled the relationships between the color metric and protein content or mitochondrial abundance (Table 1). Protein content was best described by an exponential model and showed consistent relationships across muscle types: darker muscles exhibited higher protein content in both the DLM and DVM (Figure 4B; log–log Pearson correlation: DLM *r*^2^ = 0.55, p = 0.001; DVM *r*^2^ = 0.52, *p* = 0.004). The color metric also predicted citrate synthase activity, with darker muscles exhibiting higher CS activity, consistent with higher mitochondrial abundance in these muscles. However, the form of the positive relationship difference between the two muscle types. In the DLM, the exponential model provided the best fit (Figure 4C; log–log Pearson correlation: DLM *r*^2^ = 0.45, *p* = 0.009; Table 1). In the DVM, however, the quadratic model provided the best fit, with intermediate (pink) muscles exhibiting the highest CS activity (Figure 4C). Although the exponential model had a lower AIC value (47.83) than the quadratic model (93.44) for the DVM, it explained substantially less variance (*R*^2^ = 0.023) than the quadratic model (*R*^2^ = 0.233), indicating poor explanatory power despite nominally higher AIC support.

**Table 1.** Comparison of linear, quadratic, and exponential models for flight muscle color metric and protein content or flight muscle color metric and CS activity (a proxy of mitochondrial abundance) in the flight muscles: DLM (dorsal longitudinal muscle) and DVM (dorsal ventral muscle) (Figure 4).

| Model | R<sup>2</sup> | F | df | p | AIC |
| --- | --- | --- | --- | --- | --- |
| DLM: Protein |  |  |  |  |  |
| linear | 0.277 | 11.87 | 1,31 | 0.00166 | 60.44 |
| quadratic | 0.281 | 5.849 | 1,30 | 0.00716 | 62.76 |
| **exponential** | **0.297** | **13.12** | **1,31** | **0.00103** | **39.56** |
| DVM: Protein |  |  |  |  |  |
| linear | 0.231 | 8.102 | 1,27 | 0.00834 | 66.07 |
| quadratic | 0.233 | 3.943 | 1,26 | 0.0320 | 68.00 |
| **exponential** | **0.267** | **9.846** | **1,27** | **0.00409** | **31.00** |
| DLM: CS Activity |  |  |  |  |  |
| linear | 0.111 | 3.860 | 1,31 | 0.0585 | 106.9 |
| quadratic | 0.161 | 2.879 | 1,30 | 0.0718 | 107.0 |
| **exponential** | **0.203** | **7.895** | **1,31** | **0.00851** | **57.51** |
| DVM: CS Activity |  |  |  |  |  |
| linear | 0.016 | 0.4528 | 1,27 | 0.507 | 100.3 |
| **quadratic** | **0.274** | **4.915** | **1,26** | **0.0155** | **93.44** |
| exponential | 0.023 | 0.6427 | 1,27 | 0.430 | 47.83 |

### Muscle color metric is correlated to iron level

Finally, we evaluated whether the color metric predicted iron content, an additional indicator of muscle functional state. As flight muscles are highly enriched in mitochondria, an iron-dense organelle essential for energy production in metabolically active tissues, we reasoned that iron content would likely contribute to muscle color so that muscles containing more iron would appear darker. As in prior analyses, we restricted this assessment to long-winged females and generated a second, independent dataset comprising of 38 long-winged females with muscle colors spanning red, pink, and white categories. Consistent with both the training dataset and the first test dataset, the color metric captured variation both within and among color categories, further validating its ability to quantify muscle color (Figure 5A).

**Figure 5.**
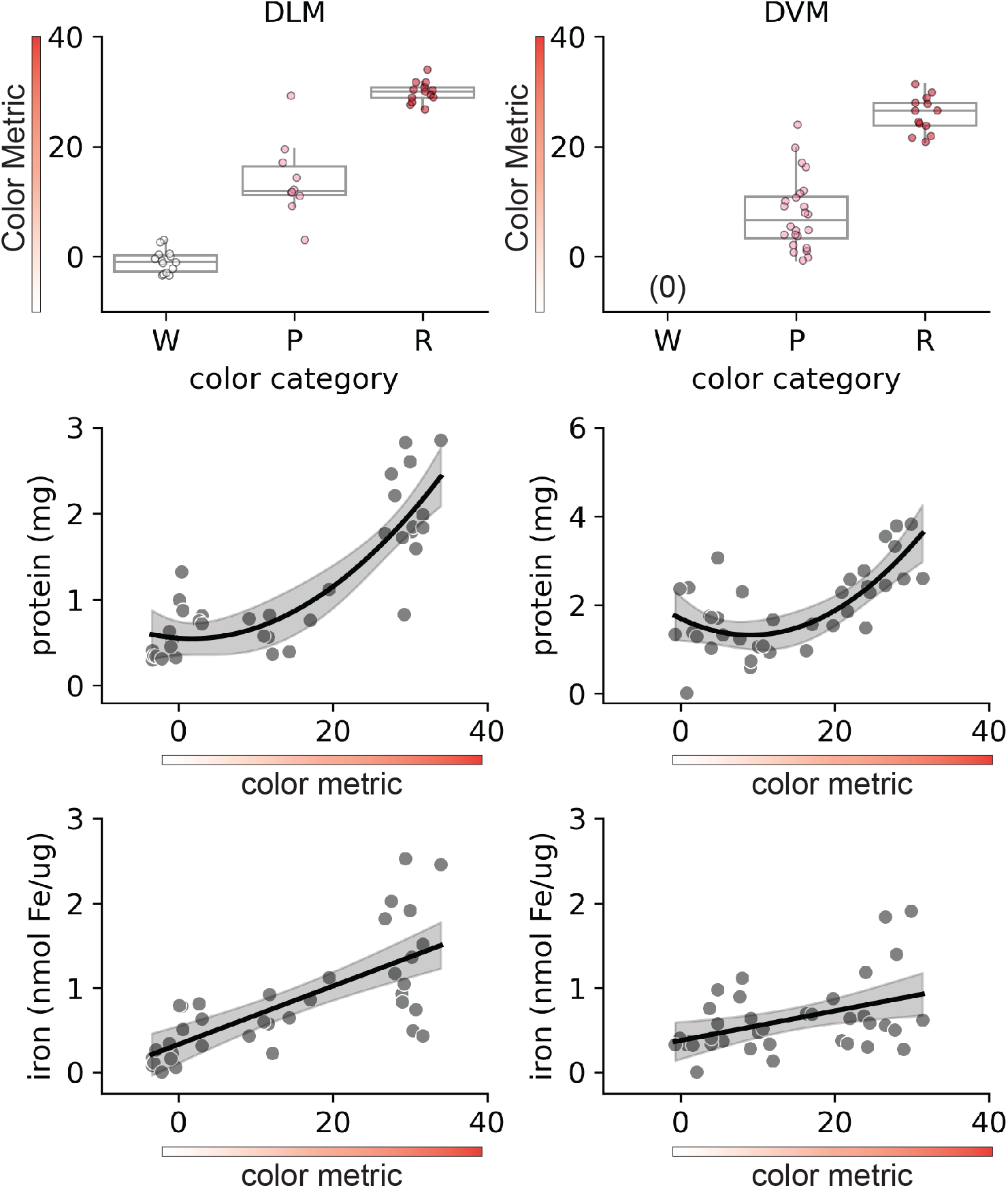
Muscle color is associated with protein content and iron in both the DLM and DVM. A) Color Metric across manual color categories (R = red, P = pink, and W = white) in the DLM (left) and DVM (right). Raw data is plotted as points, boxplots shows the quartiles, with the lower and upper edges of the box corresponding to the 25th and 75th percentile, and the whiskers indicate the values within +/-1.5 interquartile range (Tukey’s convention) to help identify extreme outliers. Scatterplots showing the relationship between Color Metric and (B) protein content and (C) iron content in the DLM (left) and DVM (right). Solid lines represent the fitted quadratic or linear model with shaded 95% confidence intervals.

Because the two test datasets comprised females of different ages (see Methods), we first modeled the relationship between the color metric and protein content to establish continuity between datasets (Table 2). Protein content was best described by a quadratic model and was consistent across muscle types, with darker muscles exhibiting the highest protein content in both the DLM and DVM (Figure 5B). We then modeled the relationship between the color metric and iron content. This relationship was best described by a linear model and was consistent across muscle types, with darker muscles exhibiting higher iron content in both the DLM and DVM (Figure 5C; Pearson correlation: DLM *R*^2^ = 0.72, p < 0.001; DVM *R*^2^ = 0.43, p = 0.008).

**Table 2.** Comparison of linear, quadratic, and exponential models for flight muscle color metric and protein content or flight muscle color metric and iron content in the flight muscles: DLM (dorsal longitudinal muscle) and DVM (dorsal ventral muscle) (Figure 5).

| Model | R<sup>2</sup> | F | df | p | AIC |
| --- | --- | --- | --- | --- | --- |
| DLM: Protein |  |  |  |  |  |
| linear | 0.683 | 75.33 | 1,35 | < 0.0001 | 47.33 |
| **quadratic** | **0.745** | **49.75** | **1,34** | **< 0.0001** | **41.20** |
| exponential | 0.690 | 77.96 | 1,35 | < 0.0001 | 39.77 |
| DVM: Protein |  |  |  |  |  |
| linear | 0.373 | 20.79 | 1,35 | < 0.0001 | 83.85 |
| **quadratic** | **0.522** | **18.53** | **1,34** | **< 0.0001** | **75.83** |
| exponential | 0.201 | 8.796 | 1,35 | 0.00541 | 94.23 |
| DLM: Iron |  |  |  |  |  |
| **linear** | **0.524** | **38.49** | **1,35** | **< 0.0001** | **49.63** |
| quadratic | 0.525 | 18.77 | 1,34 | < 0.0001 | 51.55 |
| exponential | 0.443 | 27.89 | 1,35 | < 0.0001 | 102.9 |
| DVM: Iron |  |  |  |  |  |
| **linear** | **0.184** | **7.917** | **1,35** | **0.00798** | **36.38** |
| quadratic | 0.191 | 4.012 | 1,34 | 0.0273 | 38.08 |
| exponential | 0.137 | 5.548 | 1,35 | 0.0242 | 97.55 |

## DISCUSSION

Animal models of muscle remodeling provide critical insight to the cellular and physiological pathways underlying muscle resilience and remodeling. A key requirement of an effective model system is the ability to obtain rapid, low-cost, and reliable readouts of muscle state and function. The emerging model *G. lineaticeps* naturally undergoes a non-pathological form of striated muscle breakdown, making it particularly well suited for studies of muscle remodeling (Diaz et al. 2024, Treidel et al. 2023, Arevalo et al. 2025, Wiedmer et al. 2021). However, current approaches for assessing muscle state in this system are largely qualitative, representing a significant limitation due to their subjectivity and binning of a continuous trait into artificial categories, obscuring important functionally relevant variation (Diaz et al. 2024, Treidel et al. 2023, Clark et al. 2015, Zera et al. 1997).

Here, we developed and validated a muscle color metric that provides rapid and objective quantification of muscle state. This metric robustly captures variation in muscle color across remodeling stages, including the challenging transparent stage that occurs once muscle is fully histolyzed (Figure 2 and Figure 3). Validation against functional metrics demonstrated that muscle color strongly predicts protein content across muscle types and independent datasets (Figure 4), and further revealed positive relationships between muscle color, mitochondrial abundance, and iron content, although the form of these relationships was muscle- and trait-specific (Figure 4 and Figure 5). The consistent performance of this metric across multiple independent datasets indicates that it is a broadly generalizable and robust method for assessing muscle state in this model system.

The development of a continuous metric, rather than a discrete color category, is particularly well suited to this system, as it captures a fundamentally continuous biological process. Quantitative color metrics also provide finer resolution of histolysis progression and enable more detailed examination of functional traits, thereby yielding new insights into the mechanisms underlying muscle remodeling. For example, mitochondrial abundance, assessed via citrate synthase activity, exhibited an exponential relationship with color in the DLM, a muscle specialized for flight, with darker muscles showing higher mitochondrial abundance (Figure 4). This pattern is consistent with prior work demonstrating that red, flight-capable muscles have high energetic demands (Lorenz 2007, Zera et al. 1997), and that once flight ceases, these muscles no longer need to maintain metabolic function. In contrast, mitochondrial abundance and muscle color have a quadratic relationship in the DVM, a muscle that contributes to both flight and walking (Wilson 1962, Treidel et al. 2022), with mitochondrial abundance peaking at an intermediate muscle color (Figure 4C). This pattern may reveal differences in the timing of mitochondrial degradation relative to protein loss compared to the DLM, likely reflecting the selective preservation of mitochondrial function in the DVM to maintain locomotor capacity during muscle remodeling.

Iron and protein content, on the other hand, have a consistent relationship with muscle color across muscle types. Iron content exhibited a linear relationship with muscle color (Figure 5C), suggesting that iron is a primary contributor to muscle pigmentation through the stages of breakdown. Given the exponential and quadratic relationships observed with mitochondrial abundance (see above) and the linear relationship with iron content, these results also suggest that iron is allocated to multiple cellular processes beyond mitochondrial function.

Protein content exhibited an exponential relationship with muscle color in the first test dataset and quadratic in the second (Figure 4A and Figure 5A, respectively). Model selection appears to be driven primarily by protein content in paler muscles: in the second test dataset, paler muscles contained more protein than would be predicted by an exponential model. This discrepancy may reflect the differences the age ranges represented in each dataset (0-18 days vs 5 days). Regardless, both datasets show that dark muscles, prior to the initiation of the histolysis and accompanying proteolysis, exhibit little relationship between color and protein content. However, once protein degradation passes a critical threshold, changes in structural organization such as inter-fiber spacing may amplify muscle transparency, impacting optical effects, yielding an exponential or quadratic relationship between protein content and color. Notably, this process may be influenced by aging, as evidenced by the differing relationships observed across our datasets with females of different ages. Future studies could extend this framework by incorporating behavioral assays to determine the extent to which muscle color predicts functional performance, such as the capacity to sustain flight. Integrating quantitative color metrics with behavioral measures would enable direct links between muscle remodeling state and organismal performance, providing a more comprehensive understanding of how structural and physiological changes translate into functional outcomes.

Together, these results establish muscle color as a rapid, quantitative, and biologically informative proxy for muscle state during remodeling. By capturing a continuous process with a simple and objective metric, this approach overcomes the subjectivity and limited resolution of existing qualitative assessments. The strong and repeatable relationships between muscle color and key functional traits across muscle types and independent datasets underscore the robustness and generality of this method, while the muscle-specific differences revealed by these relationships provide new insight into the coordination of structural, metabolic, and micronutrient dynamics during muscle breakdown. More broadly, this framework enables scalable and standardized assessment of muscle remodeling, positioning *G. lineaticeps* as a powerful model for studying non-pathological muscle degradation and recovery and offering a transferable strategy for quantifying tissue state in other systems undergoing dynamic structural change. Ultimately, by enabling precise and high-throughput characterization of muscle remodeling dynamics, this approach has the potential to deepen our understanding of age-related muscle diseases and disorders, thereby addressing a growing public health challenge associated with aging populations worldwide.

## ACKNOWLEDGEMENTS

We thank Savanna Wang, Makenna Siebenlist, and Haley Penfield for their help with the generation of the images of the flight muscles for the training dataset. We thank Clara Medrano, Scot Morford, and Austin Hu for their help with the generation of the samples and carrying out the ferrozine assays. We thank Thomas Naef for his insights in generating standardized images of samples.

